# Volumetric 3D Printing and Melt-Electrowriting to Fabricate Implantable Reinforced Cardiac Tissue Patches

**DOI:** 10.1101/2025.03.11.642643

**Authors:** Lewis S. Jones, Hector Rodriguez Cetina Biefer, Manuel Mekkattu, Quinten Thijssen, Alessio Amicone, Anna Bock, Miriam Weisskopf, Dennis Zorndt, Debora Meier, Li Zheng, Melanie Generali, Robert Katzschmann, Omer Dzemali

## Abstract

Cardiac patches for repairing myocardial defects require mechanically stable materials that prevent bleeding and can be implanted via suturing. The current clinical standard, bovine pericardial patches (BPPs), serve this purpose but do not degrade or integrate with the myocardium, limiting their long-term effectiveness. Therefore, we have developed the Reinforced engineered Cardiac tissue Patch (RCPatch). This multimaterial patch consists of a stiffness-tuned, cardiomyocyte-infiltrated 3D metamaterial and a suturable, hydrogel-infiltrated mesh to reduce permeability and bleeding. We first designed and computationally optimized anisotropic metamaterials using a generative modelling approach and fabricated them from biodegradable poly(ε-caprolactone) (PCL) via volumetric 3D printing (VP). The metamaterial supported the infiltration of cardiomyocytes, which maintained cell viability and contractility in vitro. In a second step, we enhanced implantability and reduced blood permeability through the patch by combining a melt-electrowritten (MEW) mesh with a fibrin hydrogel. Finally, in an acute large animal trial, the RCPatch was used on an induced myocardial defect, where it withstood intraventricular blood pressure and enabled partial hemodynamic recovery. Our findings establish a scalable framework for fabricating cardiac tissue patches that integrate mechanical reinforcement with biological function, offering a surgically implantable, and potentially regenerative solution for intraventricular myocardial repair.

**Table of Contents (ToC):** 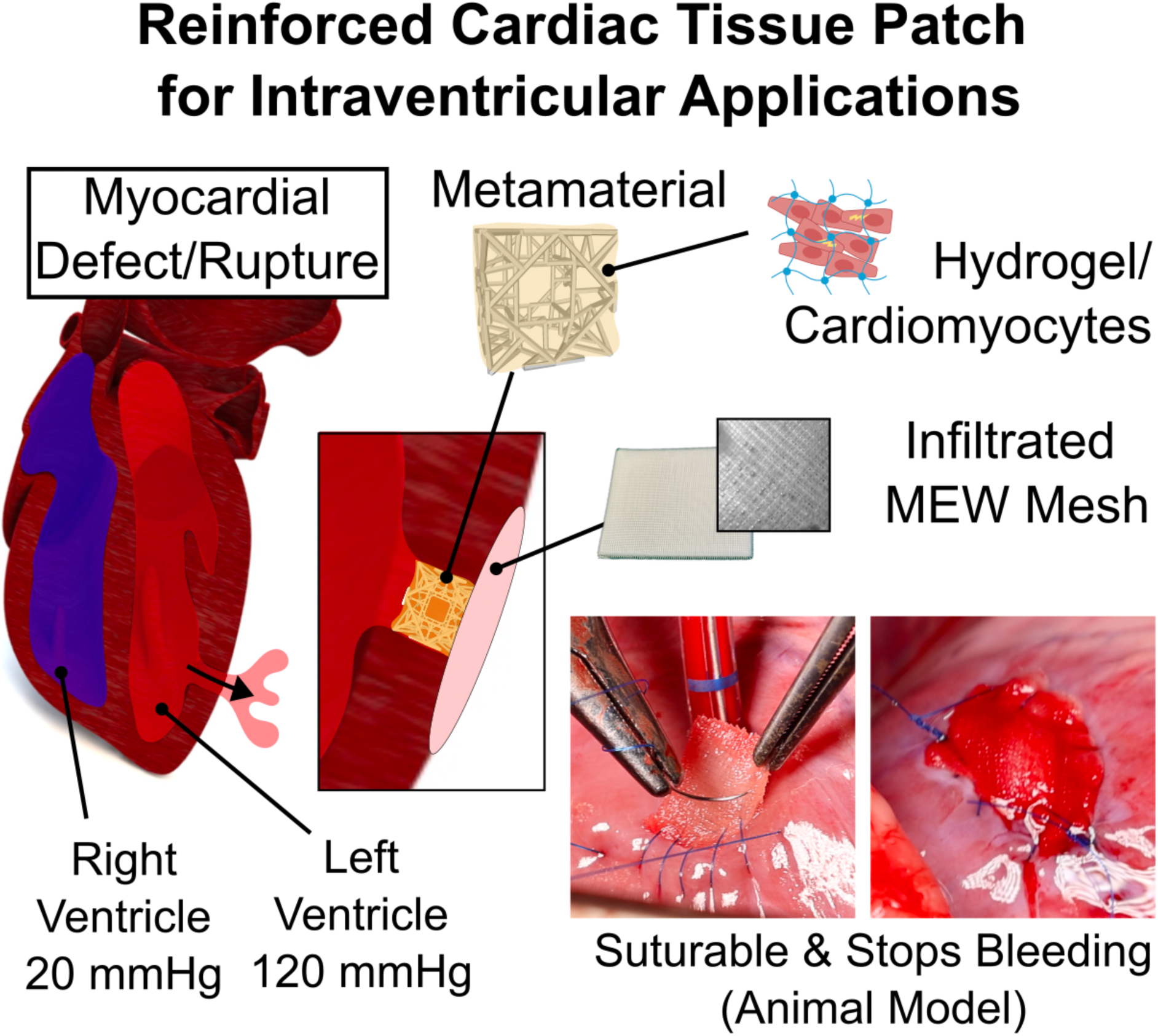

*This study presents an implantable intraventricular cardiac patch (RCPatch) combining volumetric 3D-printed metamaterials with melt-electrowritten (MEW) meshes. The design integrates tunable stiff structures with soft, cell-laden hydrogels. The RCPatch withstood suturing, intraventricular pressure, and cardiac contraction in an acute large animal myocardial defect model. The patch prevented bleeding and enabled partial hemodynamic recovery, demonstrating its potential for myocardial repair.*

## 1. Introduction

Heart disease is the leading cause of death worldwide.^[1,2]^ Myocardial Infarction (MI), a type of heart disease, occurs when blood flow to the heart is restricted, causing cardiomyocyte death, scar tissue formation, and myocardial remodeling. These changes reduce the heart’s efficiency, increasing the mechanical load on surrounding tissue and causing the infarcted region to thin.^[3]^ In severe cases, this leads to myocardial rupture, which requires immediate surgical intervention.^[4–6]^ Here, cardiac patches made from biological (bovine pericardium), or synthetic materials (polytetrafluoroethylene (PTFE), or Dacron (polyester fiber)) are implanted to re-stabilize the heart.^[7]^ However, these materials do not degrade, contract, or integrate into the myocardium.^[8]^ Furthermore, the patches undergo undesirable biological interactions such as calcification, thrombosis, and inflammation.^[9–13]^ These drawbacks hinder patch use in pediatric patients, and impair the long-term recovery and safety in many cases.^[11,14,15]^

An ideal cardiac patch would be implantable, easy to handle surgically, and provide short-term mechanical support while promoting biological regeneration of the damaged myocardium. Such a patch would fully integrate with native tissue, degrade in a controlled manner, and avoid triggering immune responses or other adverse effects. Tissue-engineered cardiac patches, or engineered heart tissues (EHTs) offer a potential solution to these challenges.^[8,16]^ Previous research has shown that large, clinically relevant cardiac tissues can be fabricated^[17]^ and engrafted onto animal hearts, where they maintain their structural and electrical properties,^[18–20]^ undergo vascularization, remuscularize, and improve cardiac function.^[21–25]^ However, tissue-engineered cardiac patches are mostly designed to be applied to the epicardial surface of the heart,^[26–28]^ and hardly any examples of intraventricular implantation exist.^[21]^ This is due to the low mechanical strength, and limited suture retainability of current tissue-engineered patches.^[8,20,29–32]^

In this work, we developed an implantable, intraventricular cardiac patch by reinforcing EHTs with 3D-printed PCL materials. A key challenge in intraventricular cardiac patch design is balancing the biological compatibility of soft materials with the mechanical robustness required for implantation. Hydrogels lack the structural integrity to withstand surgical handling and myocardial forces, while stiff synthetic materials do not support cellular integration. To address this, we used Volumetric 3D Printing (VP)^[33,34]^ to fabricate a porous PCL metamaterial which can be infiltrated with a hydrogel while providing tunable mechanical properties to match the myocardium. We combined our metamaterial with a hydrogel infiltrated melt-electrowritten (MEW) mesh, which reduces permeability and enables patch implantation via suturing. This multimaterial design allowed the RCPatch to be implanted in an acute large animal trial, where it withstood intraventricular pressure, prevented bleeding, and enabled partial hemodynamic recovery, demonstrating its potential for myocardial defect repair.

This work advances the state of the art in three key ways: (i) Moving from 2D to 3D cardiac patches by developing a mechanically reinforced 3D patch for intraventricular applications; (ii) Integrating tunable stiff materials with soft, cell-laden hydrogels to create a structurally stable yet biologically compatible cardiac patch; (iii) Using MEW meshes to enable suture retention, while hydrogel infiltration reduces permeability, preventing bleeding during implantation.

## 2. Results

### 2.1 Design of an Intraventricular Cardiac Patch

We aim to improve the current clinical standard (BPPs) by repairing myocardial defects using biodegradable and regenerative cardiac patches (Figure 1).^[11]^ Our patch, therefore, must mechanically stabilize the myocardial defect, withstand intraventricular blood pressure, and provide a source of cardiomyocytes. Designing such a cardiac patch is challenging due to the contraction of the heart and blood pressure (20-120 mmHg). Additionally, the patch must also withstand tensile forces during suturing and implantation. The patch should also encourage favorable biological processes (e.g., cellular infiltration, viability, adhesion, etc.) while resisting unfavorable ones (e.g., calcification, inflammation, hemolysis, etc.) (see Table S1, Supporting Information (SI) for a complete list of functional requirements).^[8,35]^

**Figure 1:**
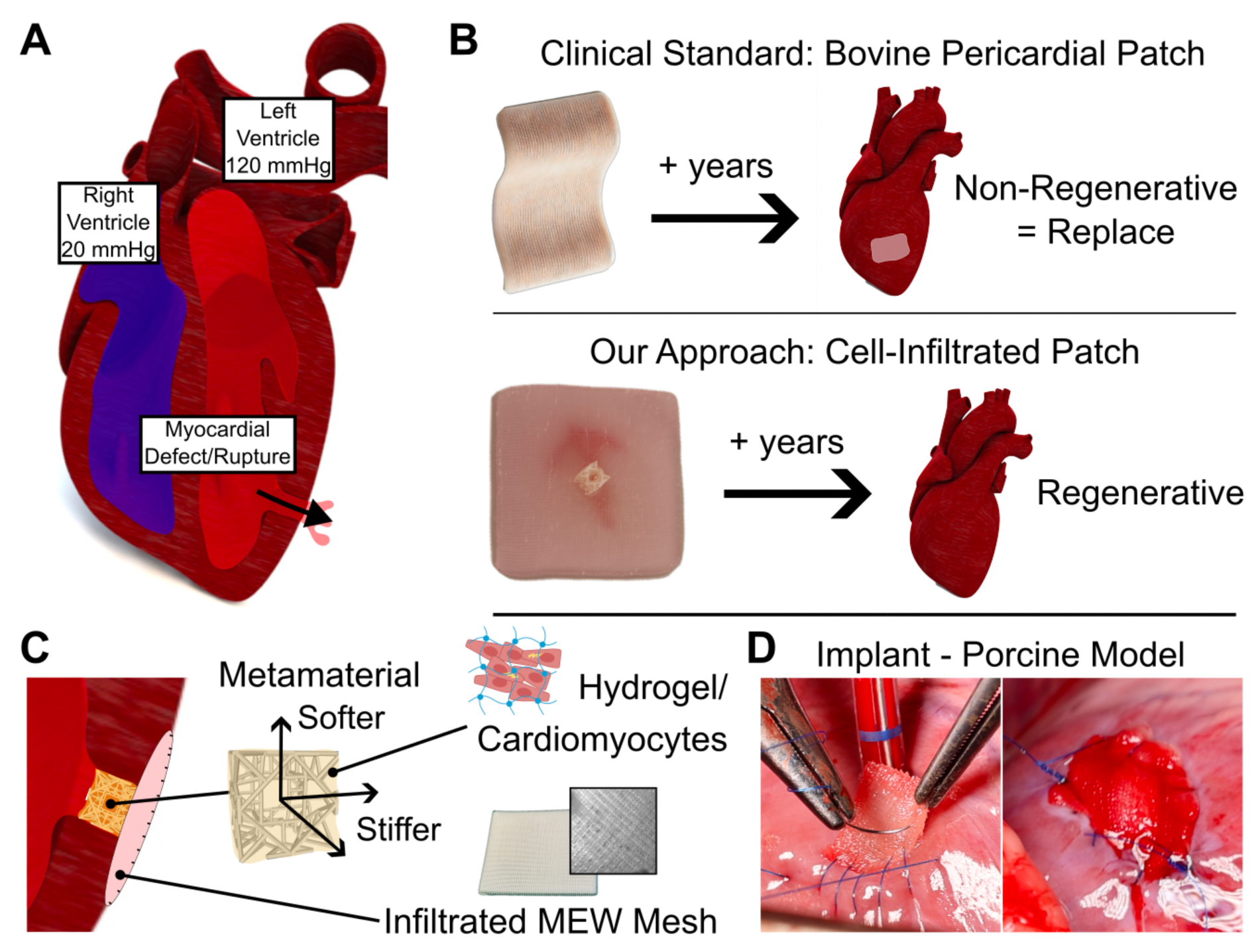
Overview of our approach to designing implantable reinforced cardiac tissue patches for intraventricular applications. (A) Schematic showing a myocardial defect. Such defects cause bleeding from the left ventricle into the pericardium (Ventricle Rupture), or the right ventricle (Septal Rupture). (B) Bovine pericardial patches (BPPs) compared with reinforced cardiac tissue patch (RCPatch). BPPs remain as permanent, non-resorbable implants, which can lead to various adverse effects. With our RCPatch, we aim for long-term integration and regeneration of the myocardium. (C) RCPatch overview. The RCPatch was fabricated by infiltrating a 3D-printed scaffold with cardiomyocytes. The scaffold comprises a metamaterial for myocardium-like material properties, and melt-electrowritten (MEW) mesh for implantability. (D) Overview of RCPatch implantation into an animal model with an induced myocardial defect (a Ø 8mm hole through the left ventricle). The images show suturing (left) and complete defect closure (right).

In our proof-of-concept study, we prioritized engineering a patch that could be implanted to restore hemodynamic stability and has the potential to be biodegradable and regenerative (Figures 1A and 1B).^[36]^ To achieve this, our RCPatch comprises three main components (Figure 1C): a MEW mesh that obstructs blood flow and allows for suturing, a VP metamaterial for cell infiltration and myocardium-like properties (Figure 1B), and iPSC-CMs to support cardiac recovery.

### 2.2. Metamaterials for Cardiac Tissue Engineering

Our cardiac patch uses a 3D-printed metamaterial to reinforce the EHT. Truss-based metamaterials were selected for their tunable and porous structure, making them well-suited for tissue engineering.^[37]^ The myocardium exhibits dynamic mechanical properties, which vary during the contraction cycle (systole/diastole) and with disease progression. Consequently, the reported tissue stiffness ranges from 20 to 500 kPa, with anisotropic and auxetic characteristics.^[38–40]^ Therefore, using metamaterials provides a highly adaptable framework, enabling precise tuning of material properties to match the specific mechanical demands of the application.

We evaluated several 3D printing techniques to fabricate high-resolution (≈ 200 µm beam diameter) truss-based metamaterials (Table S2 and Figure S1, SI). Ultimately, we decided to use Volumetric 3D Printing using photo-cross-linkable PCL (VP-PCL) (Figure 2A). This method has previously been used to fabricate structures at high resolution (≈ 100 µm beam diameters).^[33,34]^ Furthermore, PCL has various favorable physical (e.g., toughness), chemical (e.g., biodegradability), and biological properties (e.g., biocompatibility), for our application.^[21]^

**Figure 2:**
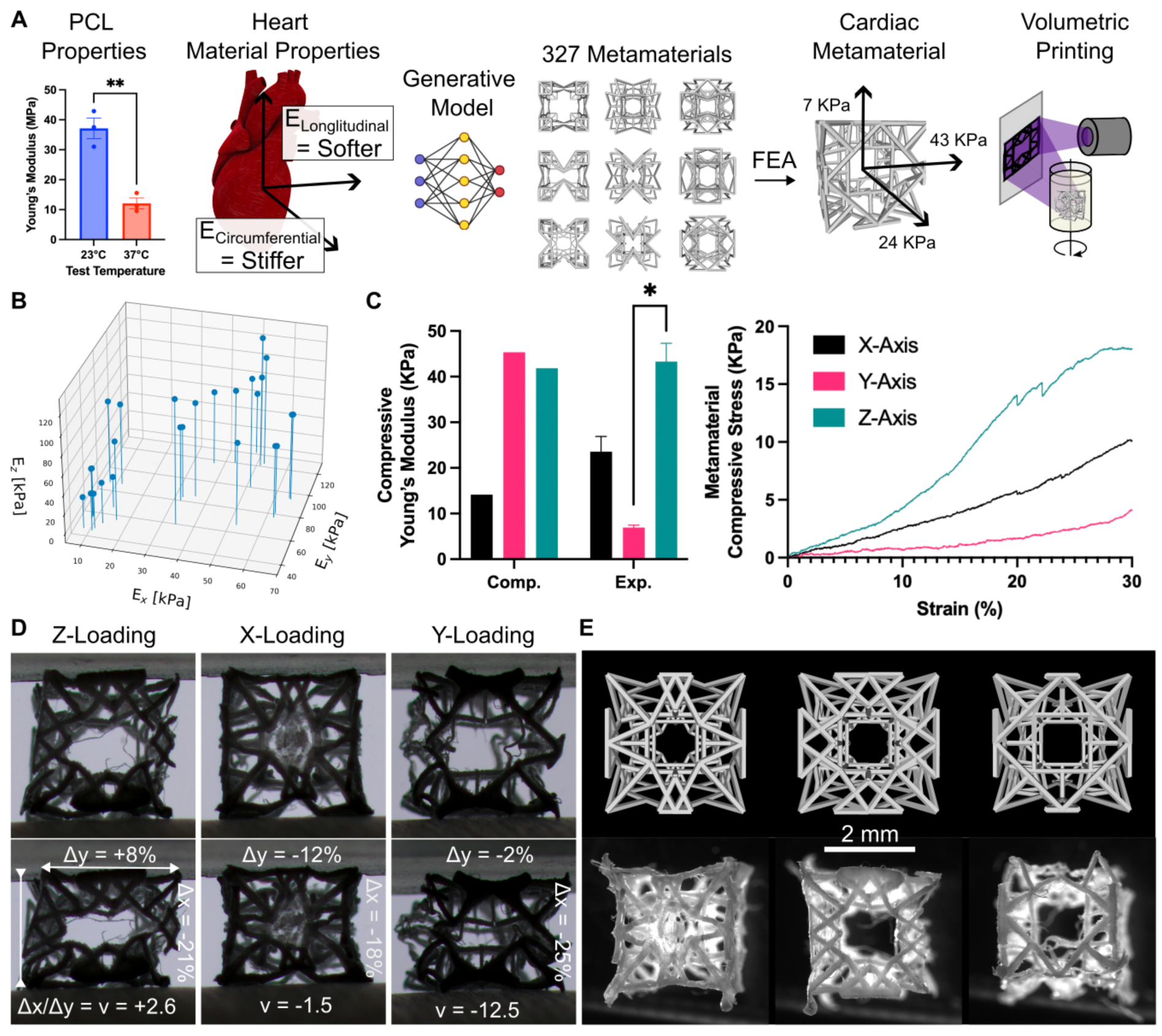
Design, Fabrication, and Characterization of Cardiac Metamaterials. (A) Cardiac metamaterial design and fabrication. VP-PCL (Volumetrically 3D Printed-PCL) material properties were obtained at 37°C (n=3, Unpaired T-Test) and used with the myocardium stiffnesses in a generative model for metamaterial structures. The stiffnesses of the resulting structures were computed with FEA. The metamaterials were then printed with VP. **(B)** Computed material properties of selected metamaterials under compression. **(C)** Left: Computational versus experimental material properties for a selected metamaterial (n = 3, one-way ANOVA, *p < 0.05). Right: experimental stress-strain curve of selected metamaterial. **(D)** Photos showing selected metamaterial under compressive load. Annotations show the change in length and width, and calculated Poisson ratio (n) under compression (≈ 20%). **(E)** Comparison between metamaterial model (top) and printed metamaterial (bottom).

We created a library of anisotropic truss-metamaterials with the desired directional Young’s moduli (*E_x_*, *E_y_*, and *E_z_*) by using a graph-based deep-learning framework^[37]^. To do this, we first measured the material properties of VP-PCL at body temperature (38 °C) (12.1 ± 1.8°C MPa, versus 37.2 ± 3.4 MPa at room temperature) (Figure 2A, Figure S2). We then used these properties, alongside the target properties of the native myocardium (one softer axis (E_Longitudinal_ ≈ 100 KPa), two stiffer axes (E_Circumferential_ ≈ 20 KPa) and auxetic) to design our truss-metamaterials. This process yielded 327 truss-metamaterials with similar mechanical behavior. These metamaterials were first evaluated using a Finite Element Analysis (FEA) model, and then according to manufacturability, to select the optimal design. The additional FE modelling was necessary due to the deviation between the simulation setup in the generative modeling framework (based on FE homogenization with periodic boundary conditions) and our single unit cell metamaterial structure.^[37,41]^

The FEA validation process yielded a sub-library of 26 metamaterials (Figure 2B), with directional Young’s moduli ranging from 9-127 KPa (X-axis (9-66 KPa), Y-axis (38-124 KPa), Z-axis (32-127 KPa)). The other truss designs were excluded due to insufficient contraction capabilities or issues encountered during the FEA simulations, such as beam collision or numerical instability due to excessive element deformation.

We selected four metamaterials from our sub-library of 26 for fabrication (Figure S3, SI) and, ultimately, one metamaterial design for mechanical characterization. This selection was based on the computed material properties and our success in printing and post-processing. We observed corresponding mechanical properties between the calculated and measured stiffness for the X-axis (14.12 KPa vs. 23.6 KPa) and the Z-axis (41.2 KPa vs. 43.3 KPa), but not for the Y-axis (45.4 KPa vs. 6.9 KPa) (Figure 2C). Additionally, while compressing the metamaterial, we noted an auxetic response (Figure 2D). The variance between the computational and experimental results is likely due to discrepancies between the metamaterial design and the printed structure (Figure 2E). For instance, certain beams did not print correctly (warped or broken), while others were over-polymerized (Figure S4, SI).

### 2.3. Cardiac Tissue Engineering using 3D Printed PCL Scaffolds

Our next objective was to engineer reinforced cardiac tissues using our 3D-printed metamaterial scaffolds (Figure 3). We infiltrated PCL metamaterials with cardiomyocytes (100 × 10^6^ cells mL⁻¹) in a fibrin hydrogel, which was injected into the scaffold (Figure 3A). To ensure consistent and well-distributed infiltration, we inserted an array of needles into the void spaces of the metamaterial. These needles provided additional wetting surfaces, stabilizing tissue formation during compaction and contraction. Additionally, the channels improved tissue perfusion, reducing cell hypoxia (Figure S5). The resulting EHTs were contractile within ≈ 3 days and showed contractility corresponding to the material stiffness (softer axis) (Video S1, SI).

**Figure 3:**
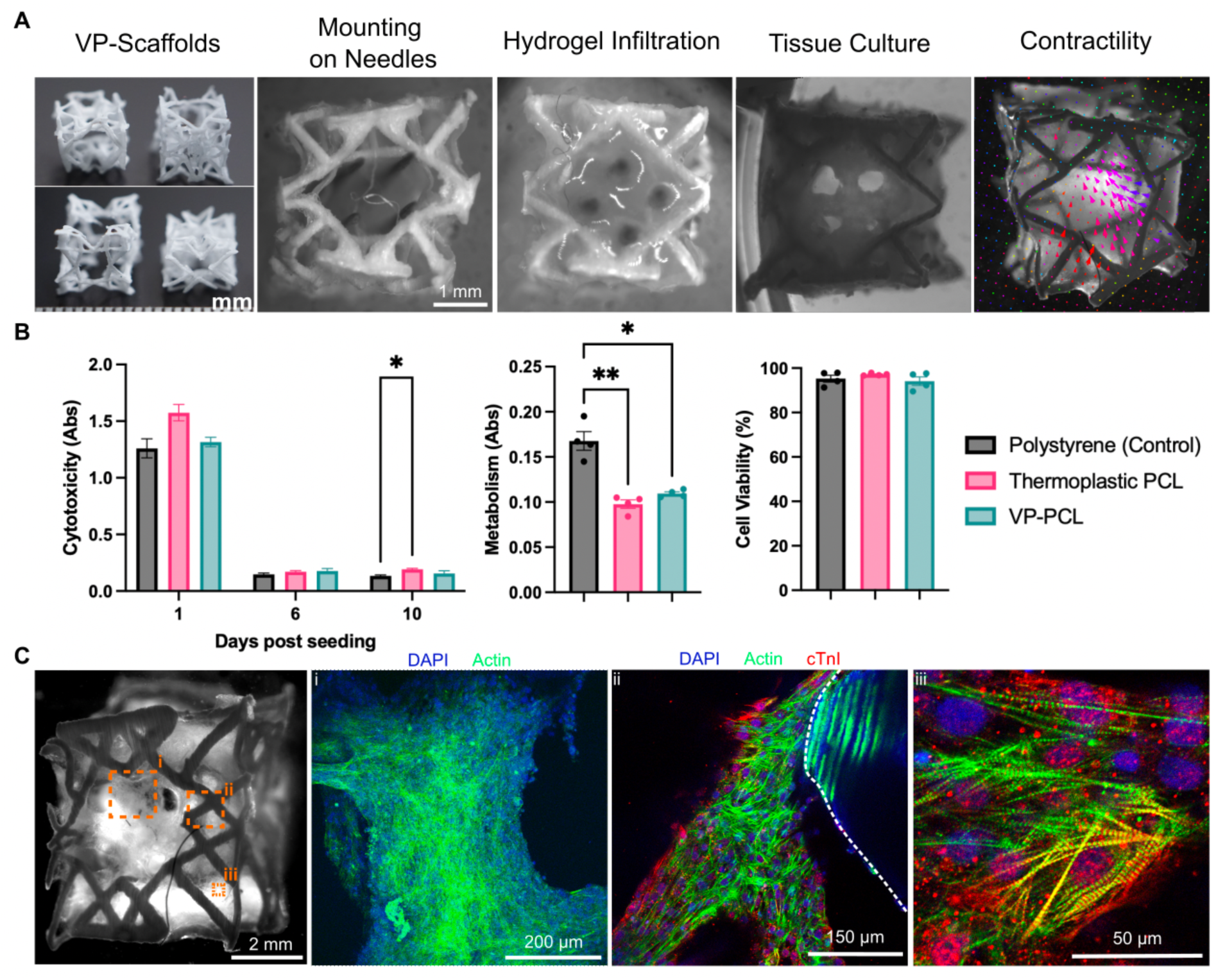
Biofabrication, Biocompatibility, and Structural Characterization of Reinforced Cardiac Tissue Patches. (A) Biofabrication of reinforced cardiac tissue patches. Cardiac Metamaterials are infiltrated with cardiomyocytes in a fibrin hydrogel and cultured, resulting in contractile cardiac tissues. (B) Biocompatibility assays of iPSC-CMs on different polymer substrates: cytotoxicity/LDH assay (left) (n=5, Two-way ANOVA), metabolism/MTT assay (middle) (n=4, one-way ANOVA) and cell viability (right) (n = 4, one-way ANOVA, *p < 0.05 and **p < 0.01). (C) Immunofluorescence staining of cardiomyocytes in infiltrated metamaterials. In panel 3, a dotted white line annotation highlights the separation between the PCL and cardiac tissue/cells.

We performed an in-depth cytocompatibility, metabolism, and viability analysis to evaluate cardiomyocytes’ compatibility with VP-PCL (Figure 3B). Specifically, we assessed cytotoxicity, metabolism, and cell viability by seeding cells in a hydrogel on various polymer substrates (Figure 2A-C). Cells seeded on VP-PCL were compared to thermoplastic PCL and polystyrene (Tissue Culture Plastic) controls. Our cytotoxicity assay revealed no significant differences among VP-PCL, thermoplastic PCL, and polystyrene across Days 1, 6, and 10. Cell metabolism was lower on both PCL substrates than polystyrene, but cell viability remained comparable across all polymer substrates. These findings suggest that VP-PCL has a similar cytocompatibility profile for cardiomyocytes to polystyrene (tissue culture plastic), a widely used substrate for cell culture.

We performed immunofluorescence staining to evaluate cardiomyocyte and EHT morphology (Figure 3C). Our EHTs contain bundles of aligned cardiomyocytes that adhere to the PCL metamaterial at multiple locations. The cardiomyocytes appear healthy and elongated, with localized sarcomere alignment influenced by local mechanical strain. Some cells exhibit sarcomere striation, confirming their structural organization. However, no global alignment direction was observed, as no external alignment methodology was applied.

### 2.4. Using Reinforced Cardiac Metamaterials for Implantation

To successfully implant the RCPatch into a ventricular defect, we needed to overcome two key challenges: reducing the permeability of the metamaterial to prevent bleeding and ensuring the patch could be securely sutured to the myocardium. We integrated a melt-electrowritten (MEW) mesh with the VP metamaterial scaffold to address these challenges. MEW uses an electrically charged nozzle to deposit molten PCL fibers precisely, allowing the fabrication of highly controlled micro- and nanoscale structures (Figure 4).^[42–44]^ First, we found that MEW meshes with a 0° to 90° grid pattern and 1.65 mm thickness (± 0.009 mm, SEM, n=3) maintain deformability within a considerable elastic range under expansion (Figure 4B). Second, we could infiltrate the MEW scaffold with a fibrin hydrogel to reduce the permeability (Figure 4C). We tested the result of this infiltration on an in vitro flow setup. This experiment showed that fluid flow through the mesh reduced ≈ 5-fold (2.4 to 0.5 L min^-1^) with hydrogel infiltration. Finally, preliminary ex vivo tests confirmed that the fibrous structure of the MEW scaffold provided sufficient mechanical integrity to be sutured to cardiac tissue.

**Figure 4:**
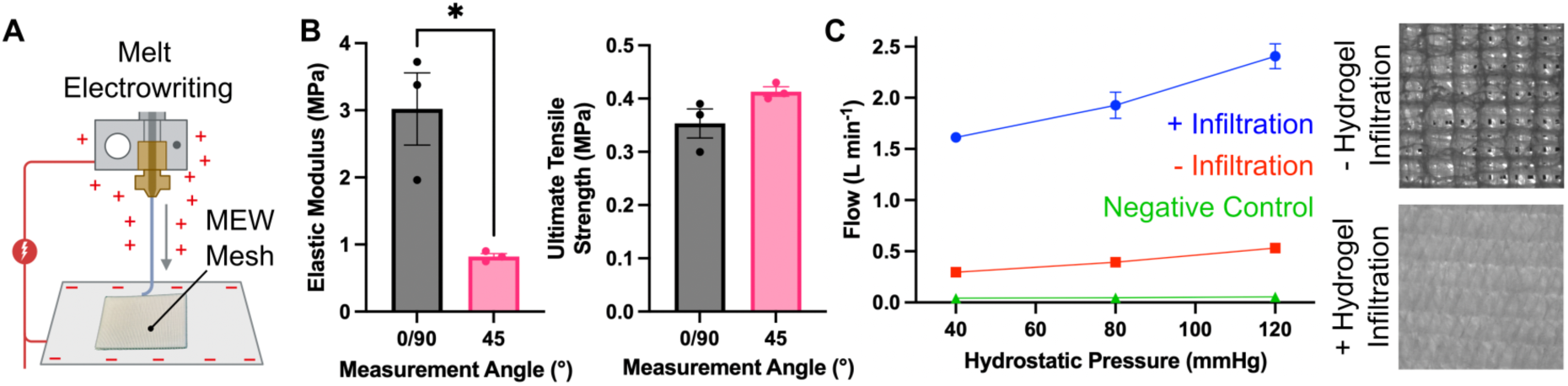
Fabrication and Properties of MEW Mesh. (A) Schematic showing mesh fabrication using melt-electrowriting (MEW). (B) Mechanical Characterization of MEW Mesh (n=3, Unpaired T-Test, *p < 0.05). (C) In vitro flow measurement of MEW mesh with and without fibrin hydrogel infiltration.

Next, we prepared for a large animal (porcine) study to assess whether our RCPatch could be sutured, withstand heart contraction and blood pressure, and restore hemodynamic stability under physiological conditions. We generated a myocardial defect model by creating a hole (⌀ 8 mm) in the left ventricle near the apex of the heart (Figure 5). Consequently, the patch must withstand the blood pressure in the left ventricle (≈ 80 mmHg).

**Figure 5:**
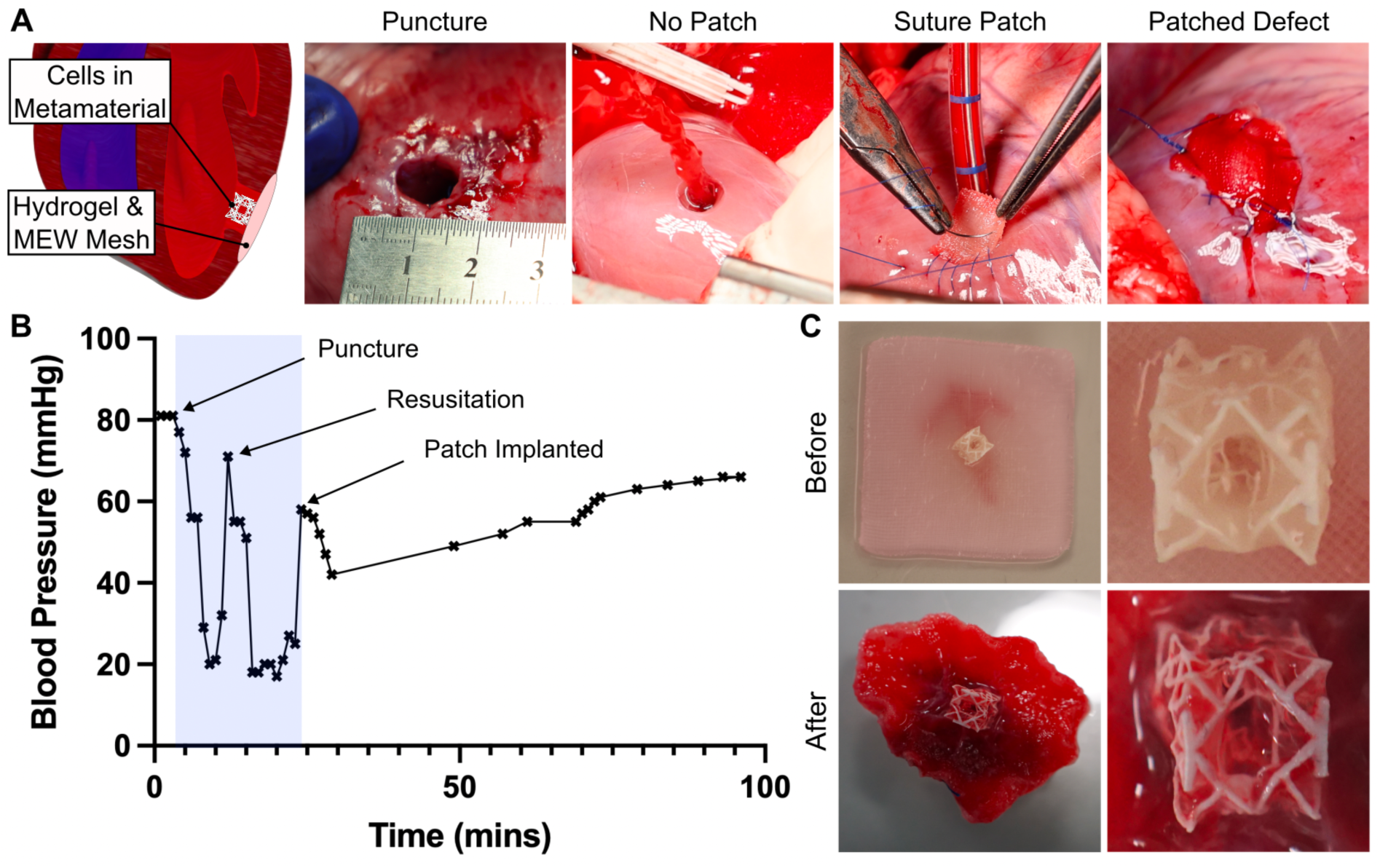
Acute Large Animal Trial using the RCPatch to repair an induced myocardial defect. (A) The RCPatch consists of a MEW scaffold combined with cardiomyocytes in a VP Metamaterial. The patch was implanted near the heart’s apex and exposed to left ventricular pressures (80 mmHg). (B) Overview of animal condition during patch implantation. The implantation procedure is highlighted in blue. (C) Images of the RCPatch before and after implantation. Note that the patch was cut to size for the implantation.

The RCPatch was implanted onto the cardiac epicardium, with the metamaterial filling the myocardial defect (Figure 5A, Video S2, SI for surgery overview). Upon implantation, most bleeding from the defect ceased immediately. Minor bleeding was observed for ≈ 10 minutes before stopping entirely. We observed partial hemodynamic recovery over the next 60 minutes, with the animal’s blood pressure increasing to 66 mmHg, compared to 80 mmHg at baseline (Figure 5B). The experiment lasted ≈ 90 minutes, where the RCPatch was implanted for ≈ 60 minutes. After termination, the patch was explanted and examined. A large amount of red blood cell infiltration was observed within the patch (Figure 5C). Furthermore, the metamaterial structure remained intact, and hydrogel and cells were observed within the metamaterial (Figure S6, SI).

## 3. Discussion

In this work, we developed an intraventricular cardiac patch (RCPatch) by combining a cardiomyocyte-infiltrated, volumetrically 3D-printed (VP) metamaterial and a melt-electrowritten (MEW) mesh. This approach integrates the properties of stiff, tunable materials with soft, cell-laden hydrogels. The metamaterial was designed through computational modeling and experimental validation, resulting in an anisotropic structure with mechanical properties matching the myocardium. The metamaterial is then infiltrated with a cardiomyocyte-laden hydrogel to form a reinforced cardiac tissue. To reduce the permeability of the patch, and enable implantation via suturing, we added the hydrogel-infiltrated MEW mesh. When implanted into an acute porcine model with an induced myocardial defect, the RCPatch withstood ≈80 mmHg, minimized bleeding, and enabled partial hemodynamic recovery. Our work extends the applicability of cardiac patches to intraventricular applications and establishes a scalable approach to tissue engineering by leveraging metamaterials, which enables the design of larger, implantable tissues for cardiac repair.

The current surgical state-of-the-art materials for cardiac tissue repair (BPPs) provide mechanical support but suffer from calcification, thrombosis, inflammation, and cannot support cell growth.^[9–13]^ Tissue-engineered alternatives, such as EHTs and hydrogels, offer a regenerative option but lack the mechanical strength to withstand suturing, blood pressure, and intraventricular contraction.^[8,20,29–32]^ Here, we used truss-based metamaterials as a porous scaffold that could be infiltrated with cells, resulting in 3D-reinforced engineered cardiac tissues. Furthermore, by using metamaterials, we could tune our material properties to the target tissue, which could enhance cell functionality and integration with the native tissue.^[37,38,45]^ In the future, we envision that this approach could be used on a larger scale, thereby providing more cells/tissue, and allowing large cardiac defects to be repaired.

The combination of VP and MEW techniques overcomes the individual limitations of each method. VP with PCL^[33]^ enables the fabrication of complex, high-resolution scaffolds with tunable mechanical properties, while the MEW mesh provides a sutureable and impermeable layer, which is critical for intraventricular applications. The woven and flexible nature of the MEW mesh is similar to surgical materials such as polyester (Dacron) and polypropylene meshes.^[7,44]^ Furthermore, the ability to print larger MEW meshes (up to 50 mm x 50 mm, determined by collector size) allows surgeons to cut the materials to size based on the defect geometry. Finally, a key success factor in the RCPatch implantability is the MEW mesh’s ability to withstand the heart pressure, and dynamic expansion/compression of the heart.^[46–48]^ In the future, the geometry of the MEW mesh could be tuned to match the properties of the myocardium, such as auxeticity.^[46]^ In addition, an integrated printing technology that combines MEW and VP could speed up patch fabrication.^[49]^

Some challenges remain: VP currently has limitations in scalability and resolution for fabricating larger metamaterials. This limitation restricted us to fabricating single-unit cell metamaterials. The resulting materials may lack the emergent behaviors which occur in larger metamaterial arrays.^[41]^ This size limitation could be addressed by optimizing the resin chemistry to minimize light attenuation^[50]^ and developing methods to increase the printable volume.^[51]^ Metamaterial fabrication using other 3D printing techniques, such as SLS, has been shown.^[52,53]^ However, this approach has a lower resolution and would, therefore, require a different type of metamaterial. Regarding the clinical translation of this research, it is important to note that we conducted only an acute animal trial (n=1). Further studies should perform chronic animal trials and evaluate the integration of the patch into the heart alongside other critical parameters such as durability, calcification, and biodegradation.

This study presents a versatile approach for designing implantable 3D cardiac tissue patches with tunable mechanical properties. Using volumetric printing, we fabricated PCL metamaterials that support cardiomyocyte infiltration. Integrating these metamaterials with hydrogel-infiltrated MEW meshes enabled suturing, reduced permeability, and prevented bleeding during implantation. In an acute large animal model, the RCPatch withstood intraventricular pressure, and enabled partial hemodynamic recovery. This approach provides a foundation for advancing the design of 3D, implantable, mechanically robust, and biological cardiac tissue patches, marking an important step toward translating engineered cardiac tissues into clinical practice for myocardial defect repair.

## 4. Materials and Methods

### Metamaterial Design and Evaluation

Our truss-metamaterial library was generated using the inverse design framework to identify truss designs that achieve the desired properties.^[37]^ We started with 500 initial guesses, selected from the training dataset based on their proximity to the target properties, and performed gradient-based optimization for each initial guess. Finally, the optimal solution was determined by FE-evaluation (COMSOL Multiphysics with a linear elastic material model, incorporating geometric nonlinearities) of the candidates. The FE model captures the influence of large deformations on the material’s mechanical response, where the relationship between stress and strain becomes dependent on changes in the structure’s geometry during loading. Using this setup, we computed the metamaterial equilibrium under a given strain. No contact model was implemented, as this would significantly reduce computational efficiency.

### Volumetric 3D Printing of Metamaterials

A photocurable PCL (VP-PCL) was prepared as previously described.^[33]^ The resin mixture contained PCL (1 g, 3.46 mmol), TPO (0.816 mg, 284 mmol), TEMPO (0.0816 mg, 19 mmol), and 4SH (32 uL, 3.5 mmol) in chloroform (214 uL). The PCL, TPO and TEMPO mixture was first sonicated and heated to 50°C, after which the 4-SH was added. The mixture was then immediately added to a glass vial and used for printing in a Tomolite (Readily 3D) using a dose of ≈ 250 mJ and the maximum attenuation constant. After printing, the mixtures were solidified by cooling, removed from the vial, washed once in chloroform, several times in acetone, and finally precipitated in ethanol and water.

### Fabrication of Melt Electrowritten (MEW) Meshes

MEW meshes were fabricated using a custom-built MEW device, using a nozzle-to-collector distance of 3.3 mm, and a polymer feed rate of 1 bar. PCL was melted at 80°C and extruded into a 0–90° grid pattern. The resulting scaffold had a pore size of 250 µm (wall spacing). The thickness of the scaffolds was measured using a 3D optical profilometer, before mechanical characterization. In preparation for the animal surgery, the VP metamaterial was attached to the MEW mesh by gluing the two structures together using the same formulation used in VP, and a UV Torch.

### Cell Culture and Hydrogel Infiltration

All materials were purchased from Gibco unless otherwise stated. Human iPSC were cultured on iMatrix-511 (Takara) coated plates in StemBrew medium (Miltenyi Biotec). The culture medium was changed every day. At 80% confluence, the iPSC were seeded for cardiomyocyte differentiation following the instructions for the StemMACS CardioDiff Kit XF (Miltenyi Biotec). iPSCs were cultured for 24h in mesoderm induction media to initiate the differentiation. Then, cells were cultured for an additional 24 h in cardiac maintenance and, afterward, cardiac induction media (24-48 h media change). Cardiomyocytes started to beat around day 7. At day 10, cells were detached using a 1:1 trypsin and Stempro accurate mixture, followed by collagenase diluted in RPME 1640 (Gibco) for cell dissociation.

A cell laden fibrin-based hydrogel was prepared by mixing fibrinogen (2 mg mL^-1^), Matrigel (10% v/v), and thrombin (50 U mL^-1^) and iPSC-CMs (≈ 100 million cells/mL) according to a previously defined protocol.^[54]^ The PCL scaffold was pre-coated with fibrinogen for 30 minutes at 37°C, and mounted upon minuten pins (2 x 2, 0.2 mm, Ento Sphinx, Czech Republic) before hydrogel infiltration. After infiltration, constructs were cultured in RPMI 1640 supplemented with B27 (2% v/v), FBS (10% v/v), Rock Inhibitor (2 μM) (StemMACS Y27632, Miltenyi Biotec), and Primocin (0.2% v/v) for 2–3 days and subsequently without the FBS and Rock Inhibitor. All cells were incubated at 37°C in an atmosphere with 95% humidity and 5% CO^2^.

### Mechanical Testing

The mechanical properties of VP-PCL were measured using an Instron ElectroPuls system with a 2527 Series DynaCell load cell (±1 kN). PCL dog bones were prepared (14 mm x 5.7 mm x 3 mm) by casting PCL into a mold and irradiating with UV. The mechanical properties of VP metamaterials were tested using the same system but using a ±250 N load cell. Compression tests were performed at a strain rate of 1 mm min^-1^ at 37°C (temperature chamber). Young’s modulus was calculated using the first 20% of the strain.

The mechanical properties of the MEW scaffolds were measured using a Zwick Roell Z005 testing machine equipped with a 5 kN load cell and a preload of 0.05 N. The test was conducted at a crosshead displacement rate of 0.1 mm/s. The specimens were rectangular, conforming to the ASTM F3510 standard, with a gauge length of 20 mm and a width of 10 mm. The MEW scaffolds’ elastic moduli were calculated per ASTM D638-22. The thickness of the samples was measured using a 3D optical profilometer (Sensofar S neox, Sensofar, Spain).

### Optical Flow Characterization

Optical Flow Analysis was conducted, as previously performed, using an open-source (OpenCV) optical flow algorithm to visualize tissue contraction.^[55]^ The contraction direction was colored using a Hue, Saturation, Value (HSV) wheel. Arrow size and opacity were scaled with the magnitude of contraction. For visibility, arrow length was scaled by a factor of 25, and only a subset of arrows were shown by uniformly sampling 1% of the pixels.

### Fluid Permeability Testing

Hydrogel-infiltrated MEW meshes were tested for permeability using an in vitro flow setup. The meshes were mounted in a hydrostatic water column, and flow was measured at 40, 80, and 120 mmHg over 30 seconds. A specialized insert was used to mount and expose the mesh (⌀6 mm) to the hydrostatic pressure.

### Cell Assays

Cells within a hydrogel were added to a 96-well plate and polymerized. Different polymer discs (thermoplastic PCL (Facilan PCL 100 Filament), VP-PCL) were then added on top of the cell/hydrogel mixture. The polystyrene control refers to tissue culture plastic, therefore no disc was added. The LDH assay (CyQUANT LDH Cytotoxicity Assay, Thermofischer) was used on the supernatant according to the manufacturer’s instructions. The MTT Assay (CyQUANT™ MTT Cell Viability Assay, Thermofischer) was performed according to the manufacturer’s instructions. An additional incubation step in Collagenase IV (5 mg mL^-1^) for 60 min was performed to release formazan crystals and ensure complete crystal dissolution. LDH and MTT Assays were evaluated on a plate reader (Tecan Infinite M200 Pro). The Live/Dead Assay (LIVE/DEAD™ Fixable Red Dead Cell Stain Kit, for 488 nm excitation) according to the manufacturer’s instructions. Before performing the assay, the tissues were fully digested in Collagenase IV (5 mg mL^-1^) for 60 min. The assay results were evaluated using Flow Cytometry (LSRFortessa, BD Biosciences).

### Large Animal Study

Animal handling procedures were performed under the Canton of Zurich guidelines for animal experimentation (ZH072/2023). The metamaterial of the RCP patch was infiltrated with cells/hydrogel 5 days before the animal surgery. On the day of the surgery, the MEW mesh was infiltrated with a fibrinogen hydrogel (2 mg mL^-1^). Animal handling procedures were performed under the Canton of Zurich guidelines for animal experimentation. The animal was anesthetized, intubated, and mechanically ventilated. A median sternotomy was performed to access the heart. A biopsy punch was used to create an ⌀ 8 mm hole through the left ventricular wall near the apex of the heart. The RCPatch was trimmed and positioned over the defect, with the VP metamaterial placed inside the ventricular wall. The MEW mesh was sutured onto the epicardium using 4-0 Polypropylene sutures. Blood pressure, ECG, and oxygen saturation were monitored continuously. Hemodynamic recovery of the animal was assessed over 60 minutes, after which the animal was sacrificed, and the patch was explanted.

### Immunofluorescence and Microscopy

Optical microscopy was performed using an inverted microscope (Olympus CKX41, with cellSens) or a stereomicroscope (Zeiss Stemi 508, with pylon by Basler). Immunostaining was performed by first washing the tissues in PBS and fixing them in 4% paraformaldehyde for 30 min at 23 °C. The tissues were permeabilized with Triton-X 100 (0.1%) in PBS for 30 min before blocking with donkey serum (5% v/v) in PBS for 2 h. Tissues were then incubated with primary antibodies (1/100, Anti-Cardiac Troponin I antibody (ab47003) in donkey serum (5%) overnight at 4 °C. Next, the tissues were washed with Tween-20 (0.1%, v/v), and incubated with secondary antibody (1/1000 Donkey Anti-Rabbit IgG H&L in donkey serum (5%) at 24 °C, for 2 h. After which an actin stain (ActinGreen 488, ReadyProbes, Thermo Fischer Scientific) was added for a further 2 h. Cell nuclei were stained in DAPI (300 μm) for 30 min at 23 °C. Samples were washed in Tween-20 (0.1%, v/v), and then imaged on a Leica SP8-AOBS-CARS Confocal Microscope.

### Statistical Analysis

Mechanical data were pre-processed to exclude artifacts caused by sample slippage, ensuring accurate Young’s modulus calculations. Data were plotted and analyzed using GraphPad Prism (x64, v. 10.4.1). Data is presented as Mean ± SEM (Standard Error of the Mean). T-test (unpaired, gaussian distribution, two-tailed) was performed in Figure 2A and 4B, one-way ANOVA (unpaired, gaussian distribution, using a Tukey test) was performed in Figure 2C and 3B (MTT and Live/Dead Assay), a Two-way ANOVA (full model with interaction; unpaired design; Sidak’s multiple comparisons; Geisser-Greenhouse correction) was performed in Figure 3B (LDH Assay). For all tests, the alpha was set to 0.05. Differences between the two experimental groups were judged to have statistical significance at \**p* < 0.05, \*\**p* < 0.01, and \*\*\**p* < 0.005; groups with no significant difference are not indicated.

## Supporting Information

Supporting Information is available from the the author.

## Acknowledgments

This work was partially funded by the University of Zurich Innopool and by a donation from Credit Suisse to the ETH Foundation to create a new chair for Robotics at ETH Zurich. RK and LJ also acknowledge funding from the SNSF Spark (CRSK-2_221397). MG acknowledges funding from the Mäxi Foundation. The authors thank the Flow Cytometry Core, Joel Chapuis for assistance with mechanical characterization, Mike Yan Michelis for assistance with Optical Flow analysis, and Marielle Airoldi for her early contributions to the project. The cardiomyocyte icon (Fig. 1C) and poly(ε-caprolactone) schematic (Fig. 4A) were created with Biorender.

## References

[1] S. S. Martin, A. W. Aday, Z. I. Almarzooq, C. A. M. Anderson, P. Arora, C. L. Avery, C. M. Baker-Smith, B. Barone Gibbs, A. Z. Beaton, A. K. Boehme, Y. Commodore-Mensah, M. E. Currie, M. S. V. Elkind, K. R. Evenson, G. Generoso, D. G. Heard, S. Hiremath, M. C. Johansen, R. Kalani, D. S. Kazi, D. Ko, J. Liu, J. W. Magnani, E. D. Michos, M. E. Mussolino, S. D. Navaneethan, N. I. Parikh, S. M. Perman, R. Poudel, M. Rezk-Hanna, G. A. Roth, N. S. Shah, M.-P. St-Onge, E. L. Thacker, C. W. Tsao, S. M. Urbut, H. G. C. Van Spall, J. H. Voeks, N.-Y. Wang, N. D. Wong, S. S. Wong, K. Yaffe, L. P. Palaniappan, on behalf of the American Heart Association Council on Epidemiology and Prevention Statistics Committee and Stroke Statistics Subcommittee, Circulation 2024, 149, e347.

[2] A. Timmis, P. Vardas, N. Townsend, A. Torbica, H. Katus, D. De Smedt, C. P. Gale, A. P. Maggioni, S. E. Petersen, R. Huculeci, D. Kazakiewicz, V. De Benito Rubio, B. Ignatiuk, Z. Raisi-Estabragh, A. Pawlak, E. Karagiannidis, R. Treskes, D. Gaita, J. F. Beltrame, A. McConnachie, I. Bardinet, I. Graham, M. Flather, P. Elliott, E. A. Mossialos, F. Weidinger, S. Achenbach, European Society of Cardiology, L. Mimoza, G. Artan, D. Aurel, M. Chettibi, N. Hammoudi, K. Vardanyan, S. Pepoyan, H. Sisakian, D. Scherr, P. Siostrzonek, B. Metzer, I. Mustafayev, T. Jahangirov, Y. Rustamova, N. Mitkovskaya, N. Shibeka, V. Stelmashok, M. De Pauw, P. Lancellotti, M. Claeys, Z. Kušljugić, A. Džubur, E. Smajić, M. Tokmakova, V. Traykov, D. Milicic, M. Pasalic, S. Pavasovic, T. Christodoulides, I. Papasavvas, C. Eftychiou, A. Linhart, M. Táborský, M. Hutyra, J. T. Sørensen, M. J. Andersen, S. D. Kristensen, M. Abdelhamid, K. Shokry, P. Kampus, M. Laine, M. Niemelä, B. Iung, A. Cohen, C. Leclercq, D. Trapaidze, K. Etsadashvili, A. Aladashvili, K. Bestehorn, S. Baldus, A. M. Zeiher, J. Kanakakis, A. Patrianakos, C. Chrysohoou, Z. Csanádi, D. Becker, Z. Járai, Þ. J. Hrafnkelsdóttir, V. Maher, J. Crowley, B. Dalton, A. Wolak, E. B. Assa, B. Zafrir, A. Murrone, C. Spaccarotella, S. Urbinati, B. Salim, S. Orazbek, A. Ayan, G. Bajraktari, D. A. Poniku, M. Erkin, A. Saamay, K. Kurban, A. Erglis, S. Jegere, I. Bajare, M. Mohammed, A. Sarkis, G. Saadeh, R. Šlapikas, T. Lapinskas, J. Čelutkienė, K. Ellafi, F. El Ghamari, J. Beissel, C. Banu, T. Felice, R. Xuereb, M. Popovici, D. Lisii, V. Rudi, A. Boskovic, M. Rabrenovic, S. Ztot, S. Abir-Khalil, J. G. Meeder, A. C. Van Rossum, M. Elsendoorn, J. Kostov, E. S. Kostovska, S. Kedev, K. Angel, O. C. Mjølstad, Ø. Bleie, M. Gierlotka, R. Dąbrowski, P. Jankowski, S. B. Baptista, J. Ferreira, V. Gil, E. Badila, D. Gaita, B. A. Popescu, E. Shlyakhto, N. Zvartau, E. Kotova, M. Foscoli, M. Zavatta, S. Stojkovic, M. Tesic, S. Juricic, G. Kaliská, R. Hatala, P. Hlivák, Z. Fras, M. Bunc, A. Pernat, Á. Cequier, M. Anguita, J. Muñiz, B. Johansson, P. Platonov, D. Carballo, M. Rüdiger-Stürchler, F. C. Tanner, H. E. Shebli, S. Kabbani, L. Abid, A. Faouzi, S. Abdessalem, V. Aytekin, I. Atar, V. Kovalenko, E. Nesukay, A. Archbold, U. Tayal, C. Wilkinson, R. Kurbanov, K. Fozilov, M. Mirmaksudov, D. Boateng, G. Daval, G. Momotyuk, D. Sebastiao, Eur. Heart J. 2022, 43, 716.

[3] K. Thygesen, J. S. Alpert, A. S. Jaffe, B. R. Chaitman, J. J. Bax, D. A. Morrow, H. D. White, The Executive Group on behalf of the Joint European Society of Cardiology (ESC)/American College of Cardiology (ACC)/American Heart Association (AHA)/World Heart Federation (WHF) Task Force for the Universal Definition of Myocardial Infarction, Circulation 2018, 138, e618.

[4] K. D. Hutchins, J. Skurnick, M. Lavenhar, G. A. Natarajan, Am. J. Forensic Med. Pathol. 2002, 23, 78.

[5] B. M. Jones, S. R. Kapadia, N. G. Smedira, M. Robich, E. M. Tuzcu, V. Menon, A. Krishnaswamy, Eur. Heart J. 2014, 35, 2060.

[6] G. J. Arnaoutakis, Y. Zhao, T. J. George, C. M. Sciortino, P. M. McCarthy, J. V. Conte, Ann. Thorac. Surg. 2012, 94, 436.

[7] M. Matteucci, D. Fina, F. Jiritano, P. Meani, W. M. Blankesteijn, G. M. Raffa, M. Kowaleski, S. Heuts, C. Beghi, J. Maessen, R. Lorusso, Eur. Heart J. Acute Cardiovasc. Care 2019, 8, 379.

[8] T. Liu, Y. Hao, Z. Zhang, H. Zhou, S. Peng, D. Zhang, K. Li, Y. Chen, M. Chen, Circulation 2024, 149, 2002.

[9] A. Salameh, W. Greimann, D. Vondrys, M. Kostelka, Semin. Thorac. Cardiovasc. Surg. 2018, 30, 54.

[10] A. A. Patukale, J. Davies, S. Marathe, N. Alphonso, P. Venugopal, World J. Pediatr. Congenit. Heart Surg. 2022, 13, 251.

[11] X. Li, Y. Guo, K. Ziegler, L. Model, S. D. D. Eghbalieh, R. Brenes, S. Kim, C. Shu, A. Dardik, Ann. Vasc. Surg. 2011, 25, 561.

[12] S. Xiao, Q. Wang, D. Fang, Z. Wang, Y. Ke, Z. Zhang, Y. Li, L. Zhong, H. Huang, Rev. Cardiovasc. Med. 2024, 25, 188.

[13] B. Poitier, J. Rancic, U. Richez, J. Piquet, S. El Batti, D. M. Smadja, Front. Cardiovasc. Med. 2023, 10, DOI 10.3389/fcvm.2023.1198020.

[14] D. F. Williams, D. Bezuidenhout, J. de Villiers, P. Human, P. Zilla, Front. Cardiovasc. Med. 2021, 8, 728577.

[15] A. D. Peivandi, S. Martens, B. Asfour, S. Martens, Pediatr. Cardiol. 2023, 44, 996.

[16] E. Rosellini, M. G. Cascone, L. Guidi, D. W. Schubert, J. A. Roether, A. R. Boccaccini, Front. Bioeng. Biotechnol. 2023, 11, DOI 10.3389/fbioe.2023.1254739.

[17] K. D. Dwyer, R. J. Kant, A. H. Soepriatna, S. M. Roser, M. C. Daley, S. A. Sabe, C. M. Xu, B.-R. Choi, F. W. Sellke, K. L. K. Coulombe, Bioengineering 2023, 10, 587.

[18] I. Y. Shadrin, B. W. Allen, Y. Qian, C. P. Jackman, A. L. Carlson, M. E. Juhas, N. Bursac, Nat. Commun. 2017, 8, 1825.

[19] C. P. Jackman, A. M. Ganapathi, H. Asfour, Y. Qian, B. W. Allen, Y. Li, N. Bursac, Biomaterials 2018, 159, 48.

[20] H. Cui, C. Liu, T. Esworthy, Y. Huang, Z. Yu, X. Zhou, H. San, S. Lee, S. Y. Hann, M. Boehm, M. Mohiuddin, J. P. Fisher, L. G. Zhang, Sci. Adv. 2020, 6, eabb5067.

[21] E. Querdel, M. Reinsch, L. Castro, D. Köse, A. Bähr, S. Reich, B. Geertz, B. Ulmer, M. Schulze, M. D. Lemoine, T. Krause, M. Lemme, J. Sani, A. Shibamiya, T. Stüdemann, M. Köhne, C. von Bibra, N. Hornaschewitz, S. Pecha, Y. Nejahsie, I. Mannhardt, T. Christ, H. Reichenspurner, A. Hansen, N. Klymiuk, M. Krane, C. Kupatt, T. Eschenhagen, F. Weinberger, Circulation 2021, 143, 1991.

[22] W.-H. Zimmermann, I. Melnychenko, G. Wasmeier, M. Didié, H. Naito, U. Nixdorff, A. Hess, L. Budinsky, K. Brune, B. Michaelis, S. Dhein, A. Schwoerer, H. Ehmke, T. Eschenhagen, Nat. Med. 2006, 12, 452.

[23] F. Weinberger, K. Breckwoldt, S. Pecha, A. Kelly, B. Geertz, J. Starbatty, T. Yorgan, K.-H. Cheng, K. Lessmann, T. Stolen, M. Scherrer-Crosbie, G. Smith, H. Reichenspurner, A. Hansen, T. Eschenhagen, Sci. Transl. Med. 2016, 8, DOI 10.1126/scitranslmed.aaf8781.

[24] R. J. Jabbour, T. J. Owen, P. Pandey, M. Reinsch, B. Wang, O. King, L. S. Couch, D. Pantou, D. S. Pitcher, R. A. Chowdhury, F. G. Pitoulis, B. S. Handa, W. Kit-Anan, F. Perbellini, R. C. Myles, D. J. Stuckey, M. Dunne, M. Shanmuganathan, N. S. Peters, F. S. Ng, F. Weinberger, C. M. Terracciano, G. L. Smith, T. Eschenhagen, S. E. Harding, JCI Insight 2021, 6, DOI 10.1172/jci.insight.144068.

[25] L. Gao, Z. R. Gregorich, W. Zhu, S. Mattapally, Y. Oduk, X. Lou, R. Kannappan, A. V. Borovjagin, G. P. Walcott, A. E. Pollard, V. G. Fast, X. Hu, S. G. Lloyd, Y. Ge, J. Zhang, Circulation 2018, 137, 1712.

[26] L. Ye, W.-H. Zimmermann, D. J. Garry, J. Zhang, Circ. Res. 2013, 113, 922.

[27] X. Mei, K. Cheng, Front. Cardiovasc. Med. 2020, 7, 610364.

[28] A. Akbarzadeh, S. Sobhani, A. Soltani Khaboushan, A.-M. Kajbafzadeh, Bioengineering 2023, 10, 106.

[29] Y. Gao, X. Han, J. Chen, Y. Pan, M. Yang, L. Lu, J. Yang, Z. Suo, T. Lu, Proc. Natl. Acad. Sci. 2021, 118, e2103457118.

[30] J. Visser, F. P. W. Melchels, J. E. Jeon, E. M. van Bussel, L. S. Kimpton, H. M. Byrne, W. J. A. Dhert, P. D. Dalton, D. W. Hutmacher, J. Malda, Nat. Commun. 2015, 6, 6933.

[31] J. Park, T. Y. Kim, Y. Kim, S. An, K. S. Kim, M. Kang, S. A. Kim, J. Kim, J. Lee, S. Cho, J. Seo, Adv. Sci. 2023, 10, 2303651.

[32] A. Pezhouman, N. B. Nguyen, M. Kay, B. Kanjilal, I. Noshadi, R. Ardehali, J. Mol. Cell. Cardiol. 2023, 182, 75.

[33] Q. Thijssen, A. Quaak, J. Toombs, E. De Vlieghere, L. Parmentier, H. Taylor, S. Van Vlierberghe, Adv. Mater. 2023, 35, 2210136.

[34] Q. Thijssen, J. Toombs, C. C. Li, H. Taylor, S. Van Vlierberghe, Prog. Polym. Sci. 2023, 147, 101755.

[35] R. Lakshmanan, U. M. Krishnan, S. Sethuraman, Expert Opin. Biol. Ther. 2012, 12, 1623.

[36] P. R. Schmitt, K. D. Dwyer, K. L. K. Coulombe, ACS Appl. Bio Mater. 2022, 5, 2461.

[37] L. Zheng, K. Karapiperis, S. Kumar, D. M. Kochmann, Nat. Commun. 2023, 14, 7563.

[38] M. Kapnisi, C. Mansfield, C. Marijon, A. G. Guex, F. Perbellini, I. Bardi, E. J. Humphrey, J. L. Puetzer, D. Mawad, D. C. Koutsogeorgis, D. J. Stuckey, C. M. Terracciano, S. E. Harding, M. M. Stevens, Adv. Funct. Mater. 2018, 28, 1800618.

[39] L. A. Reis, L. L. Y. Chiu, N. Feric, L. Fu, M. Radisic, J. Tissue Eng. Regen. Med. 2016, 10, 11.

[40] A. Silvestri, M. Boffito, S. Sartori, G. Ciardelli, Macromol. Biosci. 2013, 13, 984.

[41] R. N. Glaesener, E. A. Träff, B. Telgen, R. M. Canonica, D. M. Kochmann, Int. J. Solids Struct. 2020, 206, 101.

[42] J. C. Kade, P. D. Dalton, Adv. Healthc. Mater. 2021, 10, 2001232.

[43] A. Hrynevich, B. Ş. Elçi, J. N. Haigh, R. McMaster, A. Youssef, C. Blum, T. Blunk, G. Hochleitner, J. Groll, P. D. Dalton, Small 2018, 14, 1800232.

[44] P. D. Dalton, Curr. Opin. Biomed. Eng. 2017, 2, 49.

[45] J. Prager, C. F. Adams, A. M. Delaney, G. Chanoit, J. F. Tarlton, L.-F. Wong, D. M. Chari, N. Granger, J. Tissue Eng. 2020, 11, 2041731420934806.

[46] D. Olvera, M. Sohrabi Molina, G. Hendy, M. G. Monaghan, Adv. Funct. Mater. 2020, 30, 1909880.

[47] A. S. Federici, O. Garcia, D. J. Kelly, D. A. Hoey, Adv. Funct. Mater. 2024, 34, 2409883.

[48] Z. Dong, X. Ren, B. Jia, X. Zhang, X. Wan, Y. Wu, H. Huang, Mater. Today Bio 2024, 26, 101098.

[49] G. Größbacher, M. Bartolf-Kopp, C. Gergely, P. N. Bernal, S. Florczak, M. de Ruijter, N. G. Rodriguez, J. Groll, J. Malda, T. Jungst, R. Levato, Adv. Mater. 2023, 35, 2300756.

[50] Q. Thijssen, J. A. Carroll, F. Feist, A. Beil, H. Grützmacher, M. Wegener, S. V. Vlierberghe, C. Barner-Kowollik, Mater. Horiz. 2024, 11, 6184.

[51] A. Boniface, F. Maître, J. Madrid-Wolff, C. Moser, Light Adv. Manuf. 2023, 4, 1.

[52] J. H. Park, H.-J. Park, S. J. Tucker, S. K. Rutledge, L. Wang, M. E. Davis, S. J. Hollister, Adv. Funct. Mater. 2023, 33, 2215220.

[53] W. Y. Yeong, N. Sudarmadji, H. Y. Yu, C. K. Chua, K. F. Leong, S. S. Venkatraman, Y. C. F. Boey, L. P. Tan, Acta Biomater. 2010, 6, 2028.

[54] W. Bian, B. Liau, N. Badie, N. Bursac, Nat. Protoc. 2009, 4, 1522.

[55] L. S. Jones, M. Filippi, M. Y. Michelis, A. Balciunaite, O. Yasa, G. Aviel, M. Narciso, S. Freedrich, M. Generali, E. Tzahor, R. K. Katzschmann, Adv. Sci. 2024, 11, 2404509.

