## Supplementary Information for "Volumetric 3D Printing and Melt-Electrowriting to Fabricate Implantable Reinforced Cardiac Tissue Patches"

Other Supplementary Materials for this manuscript include:

- Video S1 – Metamaterial Contractility
- Video S2 – Overview of RCPatch Implantation

#### Table S1: Overview of the cardiac patch functional requirements, properties, results, and proposed novelty.

| Functional Requirement | Explanation | Our Approach | Property | Current Result | Novelty | Comparable References |
|---|---|---|---|---|---|---|
| **Mechanical** | | | | | | |
| Extendable/Compressable | Stretch and compress with systole/diastole. | Reinforcing PCL scaffold designs are deformable >20%. Metamaterial stiffness designed to match myocardium stiffness. | The cardiac patch can withstand acute implantation without being destroyed. | The patch was implanted during an acute animal experiment . After explantation no holes in the MEW scaffold were observed. The VP Metamaterial was slightly compressed but not destroyed. | First exaple of a 3D metamaterial tailored to a cardiac application. First example of using a MEW-enabled cardiac patch during a preclinical setting. | 1, 2 |
| High Burst Strength | Withstand pressure generated during systole (~120 mmHg). | Hydrogel is reinforced with a fine MEW mesh. | Low initial leakage rate. Any leakage is quickly blocked by blood clotting. | There was minimal leakage in the patch after implantation. The animal regained hydrodynamic stability (80 mmHg blood pressure). | Demonstration of hemodynamic stability using MEW-enabled patch. | 3 |
| Tear Resistance | Withstands suturing and other local stress | Use of woven and flexible MEW mesh using high tensile strength material (PCL) | Patch can withstand suturing and does not tear leading to blood leakage. | The patch was implanted via suturing aalong the entire circumference of the patch. No breakage was observed. | One of the first examples of suturing a MEW mesh during a preclinical trial. | 4 |
| Fatigue Resistance | Withstands repetitive mechanical loading | Not investigated | N/A | N/A | N/A | 3 |
| **Chemical** | | | | | | |
| Biodegradable | Ultimately degrades after recovery | Use of biodegradable materials (PCL) | Patch can support cardiac hemodynamic recovery and regeneration, and degrade over time. | Not investigated. However PCL is biodegradable. | N/A | 5 |
| Resistant to unfavourable biological processes (e.g., Calcification) | Material properties are not negatively affected by biological processes | Not investigated | N/A | Not investigated. | N/A | 6 |
| Enhances favourable biological processes | Supports cellular infiltration, adhesion, growth and promotes tissue recovery | Combine porous thermoplastic material with a biocompatible hydrogel; PCL is biocompatible. | Cardiomyocytes can be infiltrated into the patch; cells can adhere to patch. | Cardiomyocytes shown to be viable and functional when infiltrated into the VP Metamaterial. | Using 3D material demonstrates potential for large volume cardiac tissue engineering. | 7,8 |
| Conductive | Does not disrupt electrical signal propagation | Not investigated | N/A | N/A | N/A | |
| **Biological** | | | | | | |
| Cytocompatibility | Can support cell growth and infiltration | Use cytocompatibility materials (PCL) and 3D printing procedures (MEW and VP). | Cells can be seeded/infiltrated into patch and show normal functionality. | Various cytocompatibility markers evaluated (Mitochondrial Metabolism, LDH release, viability). All results comparable to a control. | Demonstration of compatibility between VP-PCL and Cardiomyocytes. | 6 |
| Hemocompatible | Does not damage the blood | Not investigated | N/A | N/A | N/A | N/A |
| Biocompatible | Does not adversely affect biological tissues | Not investigated | N/A | N/A | N/A | 6, 7 |
| **Other** | | | | | | |
| Customizable size and stiffness | Can be used for a range of defects. | 3D printing approach (MEW) can fabricate large patches (25 cm2). Material properties of metamaterial are tuneable. | Large patches can be produced and cut to size as required. Specific stiffnesses can be selected according to tissue type/required properties. | Current MEW scaffold size overlaps with range of available commercial patch sizes. Metamaterial stiffness is tuneable within range of native myocardium stiffness. | Demonstration of tuneable stiffness for cardiac metamaterials show applicability of metamaterial-approach for tissue engineering large volume implants. | 8 |
| Handleable/Implantable | Can be handled by surgeons and implanted by the determined methodology. | Cells are protected within 3D printed metamaterial, MEW scaffold can support suturing and handling throughout surgery. | The cardiac patch can be implanted via standard surgical procedures. | The cardiac patch was successfully implanted. | Demonstration that MEW-scaffolds are well suited for suturing and other blood-contacting applications. | 3 |

#### Table S2: Overview of possible 3D printing methods to fabricate cardiac metamaterials.

| 3D Printing Approach | Minimum Feature Size | Geometric Limitations | Material Limitations | Application Notes for Cardiac Patches |
|---|---|---|---|---|
| **Fused Deposition Modelling** | ~400 µm | Overhangs require support, minimum feature size is material dependent | Uses thermoplastic materials | Materials tend to be biocompatible, but the difficulty in printing overhangs limits the 3D complexity of the parts. |
| **SLA/DLP** | ~35-100 µm | Requires supports | Photopolymer resins | Can be used to produce detailed 3D parts, but very limited availability of cytocompatible resins. |
| **Multi-Photon Printing** | < 1 µm | Small print volume | Photopolymer resins | Print volume too small for cardiac patch application. |
| **Volumetric Printing** | ~25 µm | Limited print size (cm3), no support structures required, post processing can limit feature size | Photopolymer resins | Aimed towards bioprint+F8ng, such as with hydrogels, so cytocompatible formulations available. Print size currently limited but can be overcome in the future. |
| **Powder Bed Fusion** | ~ 100 µm | Thin features can warp, rough parts, no support structures required | Polymers, metals, and composites | Limited material selection. |
| **Vision Controlled Jetting** | 10 µm | Post-processing can limit minimum feature sizes | Photopolymer resins | Can be used to produce complex 3D parts, but no availability of cytocompatible resins. |

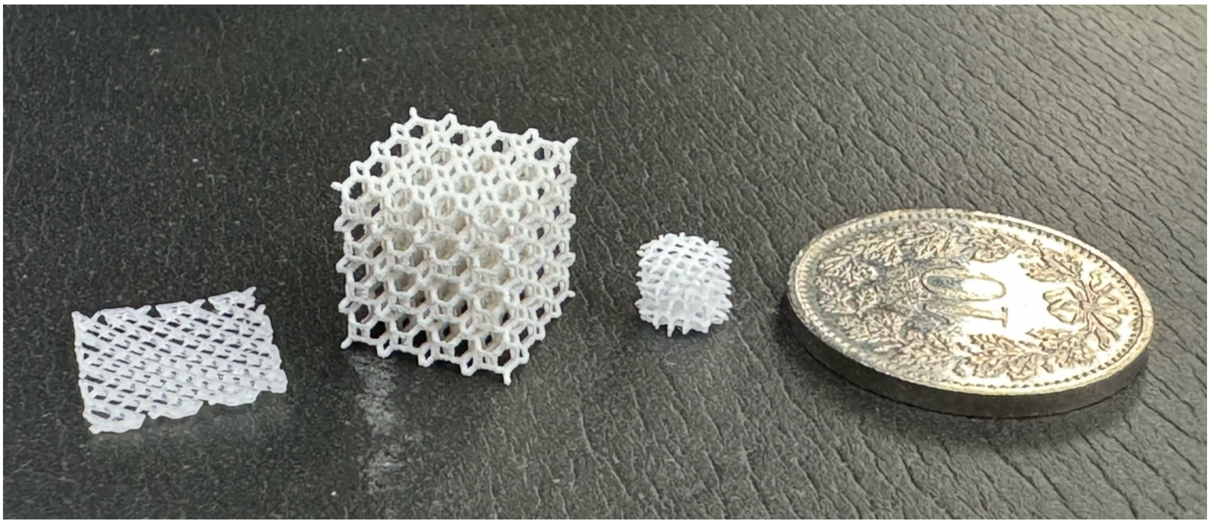

**Figure S1: Attempted Metamaterial Manufacturing Processes.** We investigated different manufacturing approaches before selecting Volumetric Printing (VP) as the preferred method. Left to right: FDM Printing, Vision Controlled Jetting (Inkbit), Volumetric Printing.

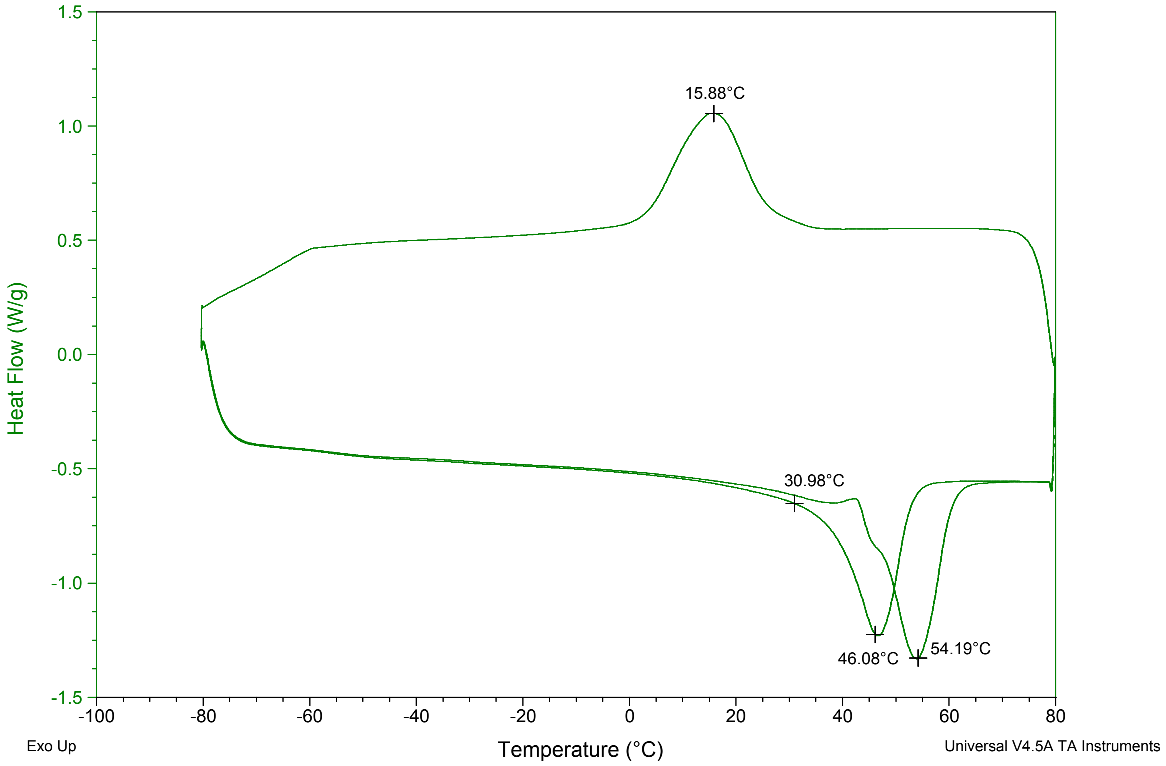


**Figure S2: Differential scanning calorimetry of VP-PCL.** The temperature transitions are labeled.

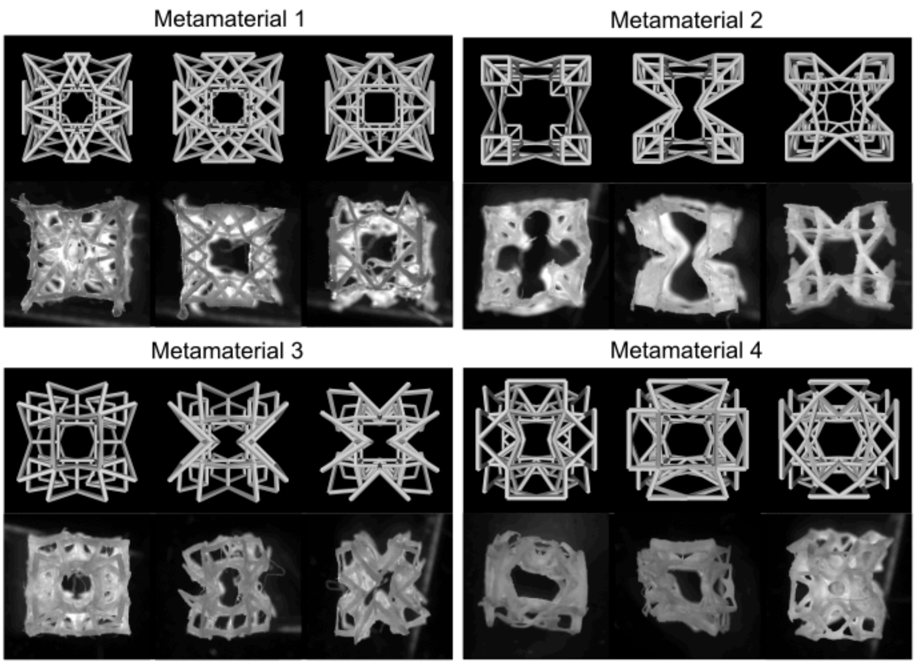


**Figure S3: Metamaterial Model/Fabrication Comparison showing four selected metamaterial geometries.**

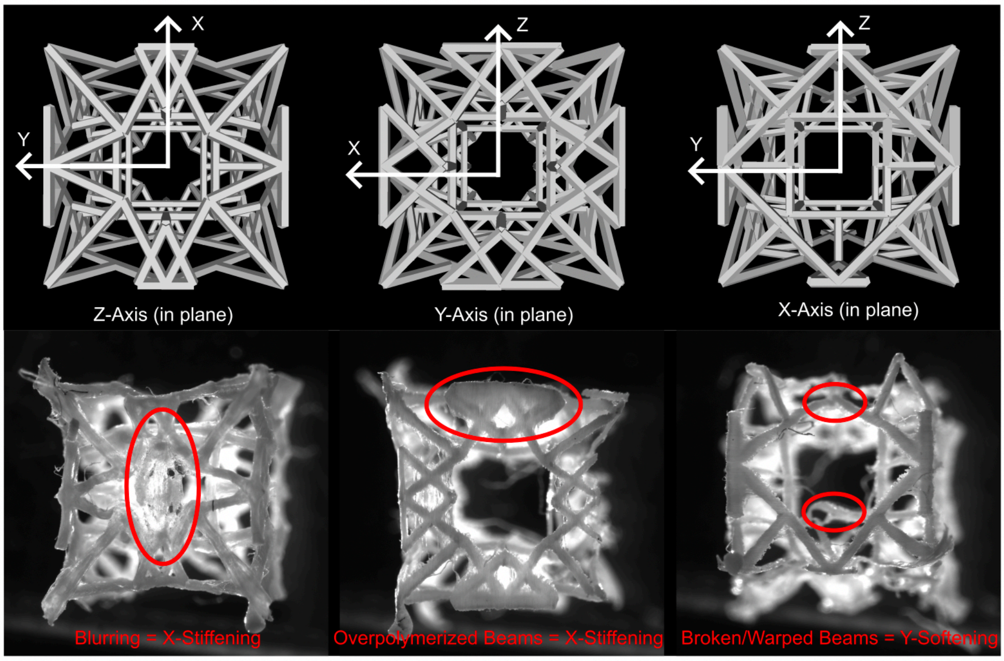


**Figure S4: Annotated Metamaterial showing regions over polymerization, blurring, and beam warping/under polymerization.**

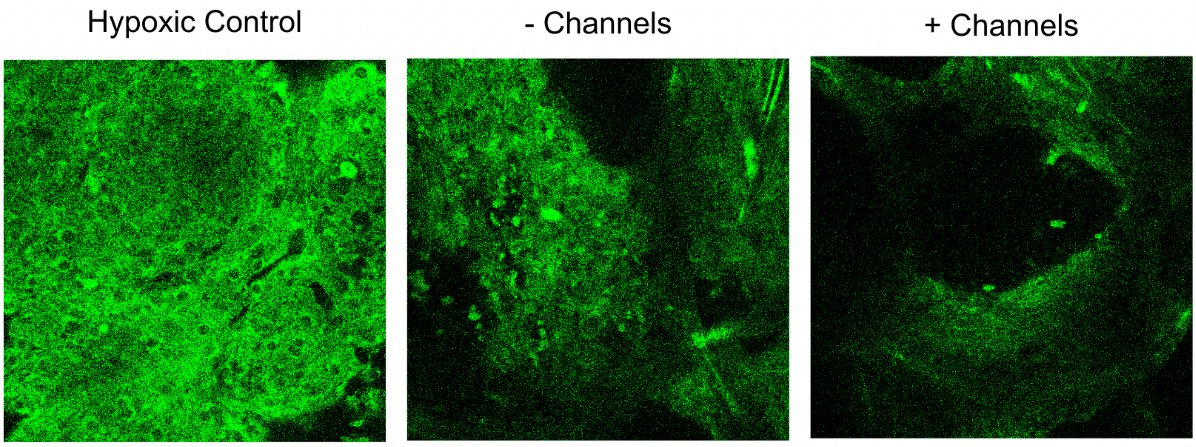


**Figure S5: Hypoxia Assay in Tissues with and without channels.** We used a fluorescently sensitive hypoxia assay, to measure cell and tissue hypoxia in tissues with and without channels. We also performed a hypoxia control (cells in 100% $CO_2$ for 4 hours).

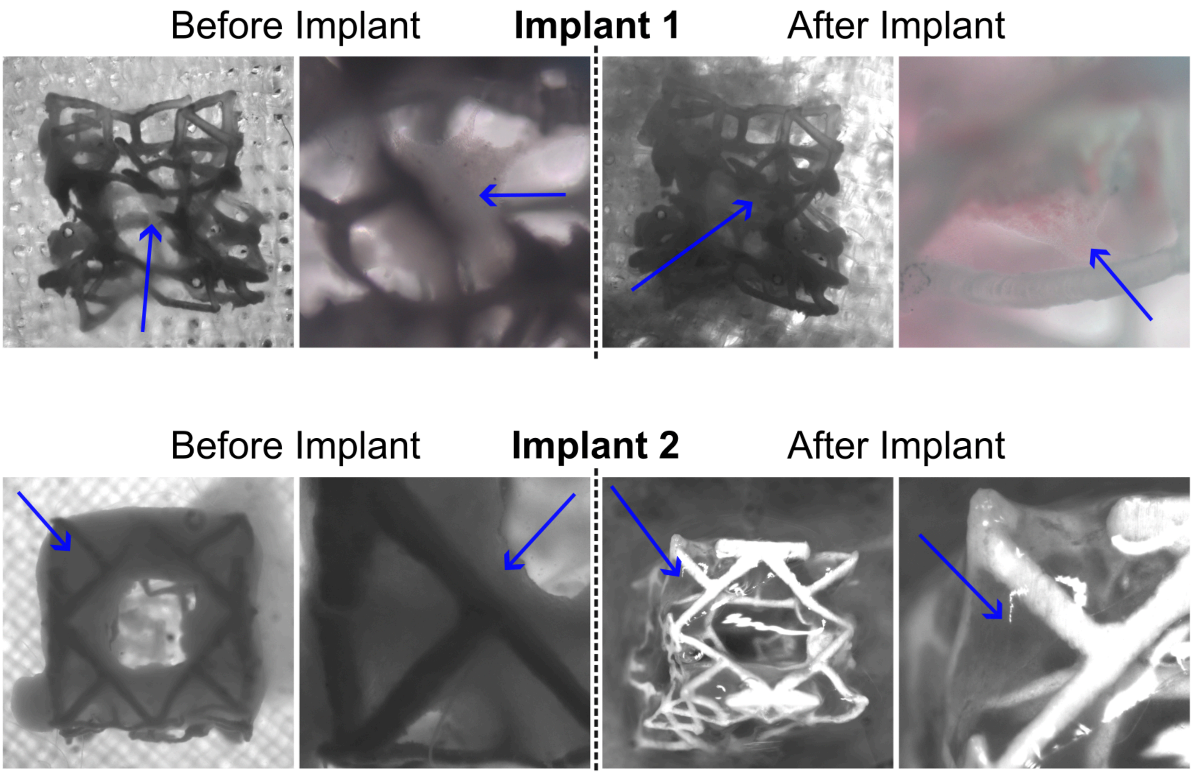


**Figure S6: Comparison of RCPatch before and after implantation.** The two different implants correspond to two different experiments. The blue arrows point to the engineered cardiac tissue contained within the metamaterial.

### References


[1] D. Olvera, M. Sohrabi Molina, G. Hendy, M. G. Monaghan, *Adv. Funct. Mater.* **2020**, *30*, 1909880.
[2] Z. Dong, X. Ren, B. Jia, X. Zhang, X. Wan, Y. Wu, H. Huang, *Mater. Today Bio* **2024**, *26*, 101098.
[3] A. S. Federici, O. Garcia, D. J. Kelly, D. A. Hoey, *Adv. Funct. Mater.* **2024**, *34*, 2409883.
[4] Y. Han, M. Lian, B. Sun, B. Jia, Q. Wu, Z. Qiao, K. Dai, *Theranostics* **2020**, *10*, 10214.
[5] Q. Thijssen, A. Quaak, J. Toombs, E. De Vlieghere, L. Parmentier, H. Taylor, S. Van Vlierberghe, *Adv. Mater.* **2023**, *35*, 2210136.
[6] P. R. Schmitt, K. D. Dwyer, K. L. K. Coulombe, *ACS Appl. Bio Mater.* **2022**, *5*, 2461.
[7] W. Y. Yeong, N. Sudarmadji, H. Y. Yu, C. K. Chua, K. F. Leong, S. S. Venkatraman, Y. C. F. Boey, L. P. Tan, *Acta Biomater.* **2010**, *6*, 2028.
[8] J. H. Park, H.-J. Park, S. J. Tucker, S. K. Rutledge, L. Wang, M. E. Davis, S. J. Hollister, *Adv. Funct. Mater.* **2023**, *33*, 2215220.
